# circVDJ-seq for T cell clonotype detection in single-cell and spatial multi-omics

**DOI:** 10.1101/2025.09.16.675546

**Authors:** Izabela Plumbom, Benedikt Obermayer, Raphael Raspe, Anna Pascual-Reguant, Ilan Theurillat, Tancredi Massimo Pentimalli, Yu-Hsin Hsieh, Marine Gil, Carola Dietrich, Michaela Seeger-Zografakis, Claudia Quedenau, Jeannine Wilde, Caroline Braeuning, Cornelius Fischer, Markus Schuelke, Volkhard Seitz, Leif S. Ludwig, Angelika Eggert, Nikolaus Rajewsky, Tatiana Borodina, Dieter Beule, Janine Altmueller, Helena Radbruch, Anja Erika Hauser, Thomas Conrad

## Abstract

Monitoring T cell repertoires in human tissues provides important insights into immune response mechanisms in cancer, infectious diseases, and autoimmunity. However, retrieving VDJ information from single-cell and spatial transcriptomics workflows with 3’-barcoding of cDNA remains resource-intensive or requires specialized sequencing equipment. Here, we introduce circVDJ-seq for simplified and cost-efficient T cell receptor (TCR) profiling from 3’-directed workflows like single-cell or single-nucleus RNA sequencing, ATAC+RNA multi-omics, and spatial transcriptomics. Application of circVDJ-seq to freshly resected neuroblastomas and postmortem lymph nodes affected by pneumonia or COVID-19 reveals distinct immune microenvironments and T cell clonality patterns, highlighting broad utility across diverse clinical contexts.

## Background

Antigen-specific T cells are key mediators of the adaptive immune response and play a central role in autoimmune disorders, cancer, and infectious disease. T cell antigen specificity and clonal identity are determined by the TCR, which is composed of either an α/β or a γ/δ heterodimer, with α/β being the most common form of TCR found on CD4+ helper and CD8+ cytotoxic T cells. The highly diverse repertoire of reactive T cells is generated by somatic recombination of the variable (V), diversity (D), and joining (J) TCR gene segments, which together define the sequence and antigen-binding specificity of the complementarity-determining region 3 (CDR3) that directly contacts the antigen presented by the Major Histocompatibility Complex (MHC) [1]. In humans, the *TCRA* locus includes approximately 50 functional Vα and 61 Jα gene segments, while the *TCRB* locus contains 52 Vβ, 2 Dβ, and 13 Jβ gene segments. During recombination, the CDR3 sequence is further diversified by random nucleotide insertions or deletions at the junctions between gene segments, leading to an estimated diversity of 10^15^ to 10^20^ possible TCR α/β combinations [2,3].

Sequencing-based TCR profiling methods can determine individual T cell repertoires at single-cell resolution and enable the monitoring of T cell clonotype dynamics in various disease settings, and advances in single-cell multi-omics analysis have enabled a more comprehensive understanding of T cell biology. Particularly, paired TCR sequencing with mRNA expression within the same cell has established a direct link between TCR repertoire and cellular phenotypes, illuminating T cell development and function [4–6]. Commonly used approaches for single-cell TCR profiling employ 5’-barcoded scRNA-seq strategies, as 3’-directed scRNA-seq approaches fail to recover the VDJ information located in the first 500 nucleotides of TCR transcripts [5]. At the same time, advanced multi-omics approaches such as paired RNA and ATAC sequencing, or spatial transcriptomics workflows such as Visium (10x Genomics) or Slide-Seq [7], are based on 3’-barcoding of transcripts and therefore lose VDJ information during fragmentation-based library preparation. Nevertheless, information on clonotypic transcriptional regulation and lymphocyte clonality within human tissues remains highly valuable as it provides important insights into adaptive immune response mechanisms and enables the identification and characterization of antigen-specific T cell clones, which can inform the development of targeted therapeutics [8].

Several recent studies have introduced modified workflows to recover VDJ sequence information from 3’-barcoded single-cell and spatial transcriptomics workflows [8–12]. However, these methods are resource-intensive as they rely on the amplification of TCRα and TCRβ sequences by multiplexed PCR with hundreds of primers that target all possible V segments, sometimes combined with prior hybridization capture of TCRα and TCRβ cDNA molecules, and often require specialized equipment for long-read sequencing. Here, we introduce circVDJ-seq, a simplified workflow for paired full-length TCR sequencing from 3’-barcoded cDNA that massively reduces the number of required gene-specific primers and is compatible with common short-read sequencing technologies. We show that circVDJ-seq efficiently recovers T cell clonotypes from 3’-barcoded cDNA libraries and is compatible with the ATAC+RNA MO and Visium assays (10x Genomics). The identified TCR sequences and clonotype frequencies are highly concordant with results obtained from the commonly used 5’-directed Immuneprofiling v2 workflow (5’IPv2, 10x Genomics). We demonstrate that circVDJ-seq can be used to spatially resolve T cell clonotypes even in challenging samples such as autopsy-derived tissue, including lymph nodes from non-COVID-19 pneumonia and COVID-19 patients. We finally reveal contrasting tumor immune microenvironments in two freshly resected high-risk and low-risk neuroblastoma patient samples (HR-NB / LR-NB), highlighting the broad applicability of circVDJ-seq in diverse clinical settings.

## Methods

### Patient recruitment and sample collection

The PBMC (n=1), lung (n=1) and lung-draining lymph node (dLN) samples (n=3) included in this study were collected and cryopreserved in the Department of Neuropathology of Charité -Universitätsmedizin Berlin as part of the COVID-19 autopsy Biobank. Donor identities were encoded at the hospital before sharing for sample processing or data analysis. All COVID-19 donors are part of the previously published cohort in [13]. All COVID-19 donors tested positive for COVID-19 in oropharyngeal swabs at the time of hospital admission.

The neuroblastoma samples (n=2) were collected and cryopreserved during surgical resections at the University Hospital Cologne as part of the neuroblastoma study NB2004/ NB2004-HR.

The colorectal cancer sample (n=1) was part of the previously published cohort in [14], and acquired during the intraoperative pathologist’s examination at Charité University Hospital Berlin. Freshly resected neuroblastoma tissue (approx. 0.1–0.4 g) was minced using scalpels, processed using the Miltenyi Human Tumor Dissociation Kit (Miltenyi, #130-095-929) and a Miltenyi gentleMACS Tissue Dissociator (Miltenyi, #130-096-427), using program 37C_h_TDK_1 for 30–45 min. Cell suspensions were filtered using 100 µm filters, pelleted by centrifugation, treated with 1 ml ACK erythrocyte lysis buffer, washed and resuspended in ice-cold PBS, and filtered using 20 µm filters. Debris was removed using the Debris Removal Solution (Miltenyi #130-109-398). Cell suspensions were analyzed to confirm cell viability > 75% using LIVE/DEAD Fixable Dead Cell Stain Kit (488 nm; Thermo Fisher) and a BD Accuri cytometer and used for single-cell library generation.

All relevant characteristics and clinical information of the donors are presented in **Additional file 1: Table S1**.

### Generation of 3’GEX single-cell RNA-seq libraries from human PBMCs and colorectal cancer cells

Single-cell capturing and library construction for fresh PBMCs and colorectal cancer cells were performed with the Chromium Next GEM Single-cell 3⍰ Reagent Kit v3.1 (10x Genomics) according to the manufacturer’s instructions. Briefly, a droplet emulsion targeting 10,000 cells was generated in a microfluidic Next GEM Chip G, with each droplet containing a single cell and 10x chemistry for cell lysis, 3’-barcoding, and reverse transcription of released mRNA. The purified cDNA was then amplified using PCR. One quarter of the cDNA library was subjected to standard library preparation and sequencing on a NovaSeq 6000 instrument (Illumina, USA).

### Generation of 5’GEX single-cell RNA-seq libraries from human PBMCs and colorectal cancer cells

Single-cell capturing and library construction were performed with the Chromium Next GEM Single-cell 5’ Kit v2 (PBMCs) or Next GEM Single-cell 5’ Kit v1 kit according to the manufacturer’s instructions. In short, a droplet emulsion targeting 10,000 cells was generated in a microfluidic Next GEM Chip K, with each droplet containing a single cell and 10x chemistry for cell lysis, 5’-barcoding and reverse transcription of released mRNA. The purified cDNA was then amplified using PCR. One quarter of the cDNA library was subjected to GEX and TCR library preparation, respectively, followed by sequencing on a NovaSeq 6000 instrument (Illumina, USA).

### Generation of single-cell Multiome ATAC+RNA libraries from human PBMCs

Single nuclei were isolated from PBMCs as described in the demonstrated protocol CG000365 Rev C (Nuclei Isolation for Single-cell Multiome ATAC + Gene Expression Sequencing, 10x Genomics). Isolated nuclei were then processed with the Chromium Single-cell Multiome ATAC + Gene Expression v1 assay (10x Genomics) according to the manufacturer’s instructions. In short, isolated nuclei were transposed in bulk solution, followed by partitioning into a droplet emulsion using a microfluidic chip, with each droplet containing a single cell and 10x chemistry enabling nuclear lysis, 3’-barcoding, and reverse transcription of released mRNA and indexing of transposed DNA. Both ATAC and gene expression (GEX) libraries were then generated from the same pool of pre-amplified transposed DNA/cDNA and subsequently sequenced on a NovaSeq 6000 instrument (Illumina, USA).

Fixed whole-cell 10× Genomics Multiome experiments were performed as previously described for DOGMA-seq [15]. Briefly, cells were fixed in 0.1 % formaldehyde for 5 min at room temperature, quenched with 0.125 M glycine, and permeabilized for 3 min on ice in lysis buffer (10 mM Tris-HCl pH 7.4, 10 mM NaCl, 3 mM MgCl_2_, 0.1 % NP-40, 1 % BSA, 1 mM DTT, 1 U µL^−1^ RNase inhibitor). Cells were then washed three times in 1 mL of wash buffer (same composition as lysis buffer but without NP-40). After centrifugation for 5 min at 500 × g, the supernatant was removed, and the cells were resuspended in 1× Diluted Nuclei Buffer (10× Genomics). Permeabilized cells were processed using the Chromium Next GEM Single-cell Multiome ATAC + Gene Expression workflow (10× Genomics, CG000338-Rev E) with the following modification. After SPRI cleanup of the pre-amplification PCR product, beads were eluted in 100 µL EB buffer instead of 140 µL. From this eluate, 25 µL were used for ATAC-seq library construction, and 35 µL were used for cDNA amplification.

### Spatial transcriptomics from human lung, lymph nodes, and neuroblastoma samples

Visualization of gene expression from fresh-frozen human lung (n=1), lymph node (n=4) and neuroblastoma tissue (n=2) was performed using 10× Visium spatial gene expression kit (10× Genomics) following the manufacturer’s protocol. Briefly, control and COVID-19 lung samples from donors categorized based on disease durations were cut into 10⍰μm sections using an MH560 cryotome (Thermo Fisher, Waltham, Massachusetts, USA), and mounted on 10x Visium slides, which were pre-cooled to −20⍰°C. The sections were fixed in pre-chilled methanol for 30 min, stained with CD45-AF647, CD31-AF594 and DAPI for 30⍰min and imaged using an LSM 880 confocal microscope (Zeiss). The sections were then permeabilized for 10⍰min and the captured mRNA was reverse transcribed and spatially 3’-barcoded followed by cDNA library amplification. Short-read sequencing libraries were constructed from one quarter of the cDNA using the 10x Genomics Visium Spatial Gene Expression 3’ Library Construction V1 Kit. Libraries were sequenced on a NovaSeq 6000 instrument (Illumina, USA). Frames around the capture area on the Visium slide were aligned manually and spots covering the tissue were manually selected based on the immunofluorescence staining, using Loupe Browser 5.1.0 software (10× Genomics). Sequencing data were mapped to the GRCh38-2020-A reference transcriptome using the Space Ranger software (version 1.3.0, 10× genomics) to derive a feature spot-barcode expression matrix.

### Generation of circVDJ-seq libraries from 3’-barcoded cDNA

Homologous ends for Gibson assembly (TCRGOT_1 and TCRGOT_12; **Additional file 1: Table S2**) were added to 1-15 ng of 3’-barcoded cDNA through PCR using the KAPA HiFi HotStart ReadyMix PCR Kit with the following temperature cycle: 95°C for 3 min, five cycles of 98°C for 20 s, 65°C for 30 s, 72°C for 2 min, and final extension at 72°C for 5 min. Amplified cDNA was purified using 0.8X AMPure XP beads and eluted in 20 μL EB. Circularization was carried out at 50°C for 1 h with NEBuilder® HiFi DNA Assembly Master Mix in a 200 μL reaction (1x CutSmart Buffer) followed by 0.9X AMPure XP bead cleanup and elution in 20 μL EB. Remaining linear DNA was removed by incubation with 6U Lambda exonuclease at 37°C for 30 min, followed by inactivation at 75°C for 10 min. Circularized cDNA was purified using 0.8X AMPure XP beads and eluted in 20 µl EB. Next, nested PCR was performed with primers targeting the 3’ untranslated region (UTR) and constant regions of TCR α and β chains (**Additional file 1: Table S2**). PCR conditions for the first amplification (12-15 cycles) were: 95°C for 3 min, 98°C for 20 s, 62°C for 30 s, and 72°C for 1 min, followed by a final extension at 72°C for 1 min. Amplified products were purified using 0.8X AMPure XP beads. A second amplification (10 cycles) used the same conditions, followed by purification and concentration with 0.9X AMPure XP beads. Successful VDJ amplification was assessed using the Agilent 4200 TapeStation High Sensitivity DNA assay and quantified by Qubit dsDNA HS assay.

2-3 ng of amplified VDJ cDNA was used to generate final circVDJ-seq libraries using the library construction kit from 10x Genomics (PN1000190). Briefly, VDJ cDNA was subjected to enzymatic fragmentation, end-repair, and A-tailing at 32°C for 2 minutes followed by heat inactivation at 65°C for 30 minutes. Subsequently, adapter ligation was carried out by incubating the sample with ligation buffer, adapter oligos, and DNA ligase at 20°C for 15 minutes. The ligated products were purified using a 0.8X SPRIselect bead cleanup. Indexed amplification was performed using indexing primers shown **in Additional file 1: Table S2**, with cycling conditions of 98°C for 45 seconds; 8 cycles of 98°C for 20 seconds, 54°C for 30 seconds, 72°C for 20 seconds; followed by a final extension at 72°C for 1 minute. Post-PCR products were cleaned with 0.9X SPRIselect beads and eluted in elution buffer. Final libraries were assessed using the Agilent 4200 TapeStation High Sensitivity DNA assay and quantified by Qubit dsDNA HS assay. In addition, libraries were quantified using qPCR and sequenced with a NextSeq 550 System Mid-Output Kit with read configuration 90–28–10-10 using custom sequencing primers indicated in **Additional file 1: Table S2**.

### Primer Design for Multiplex PCR

Primers targeting the 3’ UTR and constant regions of TCR α and β chains were designed to facilitate the detection of rearranged TCR genes post-cDNA circularization. Sequences for human *TRAC, TRBC1*, and *TRBC2* genes were obtained from the IMGT/GENE-DB database. Primer3 was used for primer selection.

### Generation of LR Spatial VDJ libraries

Long-read spatial V(D)J libraries were prepared following the protocol described by Engblom et al. [8]. Input cDNA libraries were amplified prior to target enrichment to ensure sufficient material for downstream applications. For each library, 10 µL of Visium cDNA was used and divided into subsamples containing 10 ng per PCR reaction. PCR amplification was performed using 25 µL KAPA HiFi HotStart ReadyMix (2×; Roche), 7.5 µL cDNA primers (10x Genomics, PN 2000089), and 17.5 µL sample diluted in Milli-Q water, resulting in a total reaction volume of 50 µL. PCR cycling conditions were as follows: initial denaturation at 98 °C for 3 min; 5 cycles of 98 °C for 15 s, 63 °C for 30 s, and 72 °C for 2 min; followed by a final extension at 72 °C for 1 min. PCR subsamples were subsequently pooled and purified using 0.6X SPRIselect beads (Beckman Coulter) and eluted in 40 µL elution buffer. Subsequently, hybridization capture was performed according to the protocol xGen Hybridization Capture of DNA Libraries (version 4, Integrated DNA Technologies). The optional AMPure XP Bead DNA Concentration Protocol was applied, followed by the Tube Protocol starting from step 8. The hybridization reaction was carried out overnight (16 h). xGen Universal Blockers used for the libraries were a Read1 + polyT blocking oligo (51-CTA CAC GAC GCT CTT CCG AT TN NNN NNN NNN NNN NNN NNN NNN NNN NNT TTT TTT TTT TTT TTT TTT TTT TTT TTT TTV N-3⍰) and a TSO blocking oligo (5⍰-CCC ATG TAC TCT GCG TTG ATA CCA CG CTT-3⍰). TCR discovery probe pools were combined at a final ratio of 1:3. Following hybridization capture, post-capture PCR amplification was performed using 25 µL KAPA HiFi HotStart ReadyMix (2×), 7.5 µL cDNA primers (10x Genomics, PN 2000089), and 17.5 µL enriched library from the previous step. PCR cycling conditions were: 98 °C for 3 min; 12 cycles of 98 °C for 15 s, 63 °C for 30 s, and 72 °C for 2 min; followed by a final extension at 72 °C for 1 min. PCR products were purified using 0.8× AMPure XP beads (Beckman Coulter) and eluted in 20 µL elution buffer. The enriched libraries obtained from hybridization capture were used as input (500 ng) for SMRTbell library preparation (PacBio) as in [8].

### MAS-ISO-seq

cDNA from the 10x Multiome assay was used as input for MAS-ISO-seq library preparation (Pacific Biosciences). Single-cell cDNA molecules were first depleted of template-switching oligo (TSO) artifacts, then 16x concatenated to form ordered arrays. Following MAS adapter ligation, the library was sequenced for 24 h on the PacBio Sequel II platform. Sequenced HiFi reads were segmented into individual segmented reads (S-reads) that represent the original cDNA sequences using SMRT Link Version 11.1.0.166339.

### Single-cell and spatial transcriptomics data analysis

Due to the involvement of multiple collaborating groups, single-cell, Multiome or Visium gene expression and/or chromatin accessibility data were processed with different versions of Cell Ranger or Space Ranger against the GRCh38 reference genome, as summarized in Table 1 [16].

**Table 1:** Cell Ranger versions used for analysis.

| material | sample(s) / method | program (version) | command used |
| --- | --- | --- | --- |
| CRC | 5'IPv2 | Cell Ranger v3.0.2 | cellranger multi |
|  | 3'GEX v3.1 | Cell Ranger v3.1.0 | cellranger count |
| PBMC | 5'IPv2 | Cell Ranger v6.1.2 | cellranger multi |
|  | 3'GEX v3.1 | Cell Ranger v6.1.1 | cellranger multi |
|  | MO | Cell Ranger-arc v2.0.0 | cellranger arc count |
|  | MO + PFA | Cell Ranger-arc v2.0.2 | cellranger arc count |
| neuroblastoma | HR + LR | Space Ranger v1.3.0 | spaceranger count |
| lung tissue | chronic COVID-19 | Space Ranger v1.3.1 | spaceranger count |
| lymph node | control/acute/chronic/prolonged | Space Ranger v1.3.1 | spaceranger count |

PBMC gene expression data were analyzed in R (v4.3.2) using Seurat (v4.4.0) [17]. We used all cells with less than 10% mitochondrial RNA and with at least 250 and at most 8000 genes. We used sctransform (v0.3.5) [18] and the MapQuery workflow to transfer cell type annotations from a PBMC reference [17] . We removed cells with a layer 1 cell type prediction score below 0.75, predicted monocytes, and doublets predicted by DoubletFinder [19].

Visium data for neuroblastoma, lung tissue, and lymph node samples were loaded into R, normalized with sctransform, and subjected to PCA, neighbor graph construction, clustering and UMAP reduction. Neuroblastoma datasets were merged before PCA.

5’ gene expression data for colorectal cancer (CRC) (originally published in [20]) was integrated with 3’ data (obtained from http://zenodo.org/records/10692019) using scANVI [21] as described in [22] and learned cell types as well as UMAP coordinates were extracted.

### circVDJ-seq data analysis

Technical replicates of PBMC data generated using different assays were processed individually but also jointly after pooling reads from replicates for better comparison of assays or processing tools.

#### Cell Ranger processing

We modified Cell Ranger v6.0.0 to allow for cell barcodes from other assays than 5’IP by exchanging the file Cell Ranger-6.0.0/lib/python/Cell Ranger/barcodes/737K-august-2016.txt for corresponding files in the same directory, namely 3M-february-2018.txt (single-nucleus or 3’ assay), 737K-arc-v1.txt (multi-ome assay) or visium-v1.txt (Visium V1). We further switched fastq files for read mates 1 and 2 and then ran Cell Ranger vdj using the refdata-Cell Ranger-vdj-GRCh38-alts-ensembl-5.0.0 reference provided by 10X.

#### MiXCR processing

We used MiXCR v4.1.2 [23] with the following command line mixcr -Xmx64g analyze 10x-vdj-tcr --species hsa /path/to/read_2.fastq.gz /path/to/read_1.fastq.gz output --tag-pattern “^(CELL:N{16})(UMI:N{12})\^(R2:*)” --threads 8 -M refineTagsAndSort.whitelists.CELL=“file:3M-february-2018.txt” for the 3’ assay and the corresponding barcode whitelist files for the other assays, and 10nt UMIs for 5’IPv2.

#### TRUST processing

We used TRUST4 v1.1.5 [24] with the following command line:

TRUST4/run-trust4 -f /path/to/hg38_bcrtcr.fa --ref /path/to/human_IMGT+C.fa -u /path/to/read_2.fastq.gz --barcode /path/to/read_1.fastq.gz --barcodeRange 0 15 + --barcodeWhitelist 3M-february-2018.txt --UMI /path/to/read_1.fastq.gz --umiRange 16 25 + --od output -t 8 –repseq, again using corresponding barcode whitelists for the other assays and 10nt UMIs for 5’IPv2.

For the MAS-ISO-seq data, we used the tagged, refined, sorted, and de-duplicated bam file from the extracted mRNA sequence as read mate 2, and cell barcode + UMI as read mate 1, and used TRUST4 as follows:

TRUST4/run-trust4 -f /path/to/hg38_bcrtcr.fa --ref /path/to/human_IMGT+C.fa -1 /path/to/read1.fastq.gz -2 /path/to/read2.fastq.gz --barcode /path/to/read1.fastq.gz –UMI /path/to/read1.fastq.gz --readFormat bc:0:15,um:16:27 -o output -t 8 Cell barcodes returned by TRUST for MAS-ISO-seq data were reverse-complemented.

#### Dandelion processing

We used Dandelion v0.3.7 [25] and followed the re-annotation vignette, i.e. we used ddl.pp.format_fastas(…) and then ddl.pp.reannotate_genes(.., loci=“tr”, reassign_dj=True, org=‘human’, extended=True, flavour=‘strict’, min_j_match=7, min_d_match=9, v_evalue=0.0001, d_evalue=0.001, j_evalue=0.0001, dust=‘no’, db=‘imgt’)

#### CircVDJ-seq data post-processing and filtering

For single index libraries, we implemented an additional filtering strategy to identify and remove spurious contigs with identical cell barcode and CDR3 sequences that originated from different samples sequenced on the same flow cell and likely resulted from index hopping, as demultiplexing of the first set of libraries had to be performed in single-index mode due to failure of the initial custom index 2 sequencing primer. For this, we assigned a clone as primary (defined via cell barcode and junction aa sequence) if it either contained more than 1% of the total reads (for replicates derived from the same cDNA sequenced on the same flow cell), or if it originated from the sample with the highest read count for that clone on that flow cell. After updating the index 2 primer design, we validated that filtering for index hopping can be omitted for circVDJ-seq libraries in dual index mode.

To further account for spurious signals likely created during PCR amplification, we extracted UMI sequences for all reads assigned to a clone by Cell Ranger, grouped UMIs within Hamming distance 1, and counted the number of perfectly matching reads (using the CIGAR string) as well as the fraction of reads aligning to the 3’ end of the contig. We then assigned revised UMI and read counts to a clone by counting only UMIs containing at least one perfectly matching read but with less than half of reads aligning to the 3’ end.

#### SpatialVDJ data processing

Raw reads were preprocessed as described in the SpatialVDJ methods section [8], with the modification that all reads in the input FASTQ file for MiXCR were stored in the 5’→3’ orientation of the coding strand. V(D)J assignment was performed using MiXCR v4.7.0 with a customized YAML configuration file based on the generic-pacbio-with-umi preset, incorporating the following modifications: the saveOriginalReads=true parameter was added to the align command, and clnaOutput=true was added to the assemble command, enabling subsequent export of complete reads for all alignments. Throughout the pipeline, the full 28 bp barcode sequence was treated as a single UMI; splitting into the 16 bp spatial barcode and 12 bp UMI barcode was performed in a downstream step. Alignments were exported using mixcr exportAlignments with the following fields: V gene, V hit score, D gene, J gene, C gene, C hit score, isotype, clone ID with mapping type, clone ID, read IDs, read 1 descriptors, and molecule tag.

Reads carrying TCR alignments were extracted and saved in FASTA format for reannotation using IgBLAST v1.22.0. IgBLAST was run with the following parameters: IMGT human TR germline databases for V, D, and J segments; organism set to human; IMGT domain system; sequence type TCR; and output format 19 (AIRR-compliant TSV).

For each UMI, a single best consensus read was selected to resolve cases where reads within a UMI group differed in sequence. This was performed in three steps: (1) reads were grouped by UMI; (2) if one or more productive reads were present, non-productive reads were excluded; (3) the read with the highest germline identity was selected, with sequence length used as a tiebreaker in cases of equal identity scores.

#### Data analysis and statistics

Filtered dandelion output was combined with gene expression data by matching cell barcodes, keeping only productive and primary contigs, and filtering out secondary α or β chains for single-cell (but not Visium) data.

To impute missing α or β chains, we collected the most frequently observed combination across all our PBMC or CRC data generated using different assays and replaced a missing chain with the partner from that combination.

For the co-occurrence analysis, we re-created the co-occurrence function from squidpy in R [26]. Briefly, we created a distance matrix from the tissue coordinates of all spots and then assessed the co-occurrence probability as the ratio of the probability to observe a spot of a certain type within a radius around spots of a given index type by the probability of observing spots of a certain type overall. This calculation was repeated multiple times for random subsets of 50% of the respective index spots to obtain error bars from standard deviation.

## Results

### circVDJ-seq simplifies TCR profiling from 3’-barcoded cDNA libraries

Here we present circVDJ-seq, a simple and cost-efficient workflow for TCR profiling from 3’-barcoded single-cell cDNA libraries that is independent of large oligonucleotide panels for PCR or hybridization capture and does not require prior knowledge of all possible V gene sequences or specialized sequencing equipment. cDNA is first tagged with homologous overhangs to facilitate circularization via Gibson assembly [27] and generate a cDNA library in which the VDJ region is juxtaposed to the unique molecular identifier (UMI) and cell barcode at the 3’ end of cDNA molecules (**Fig. 1A**). A VDJ library is then amplified via nested PCR using primers against the TCRα and TCRβ constant and 3’ UTR regions (**Fig. 1A; Additional file 2: Fig. S1A; Additional file 1: Table S2**). Subsequent library preparation steps, including fragmentation, end repair, A-tailing, adapter ligation, and index PCR, can be performed with the 5’IPv2/3 TCR library preparation kit (10x Genomics) and adjusted index primers (**Fig. 1A; Additional file 2: Fig. S1B; Additional file 1: Table S2**).

**Fig. 1.**
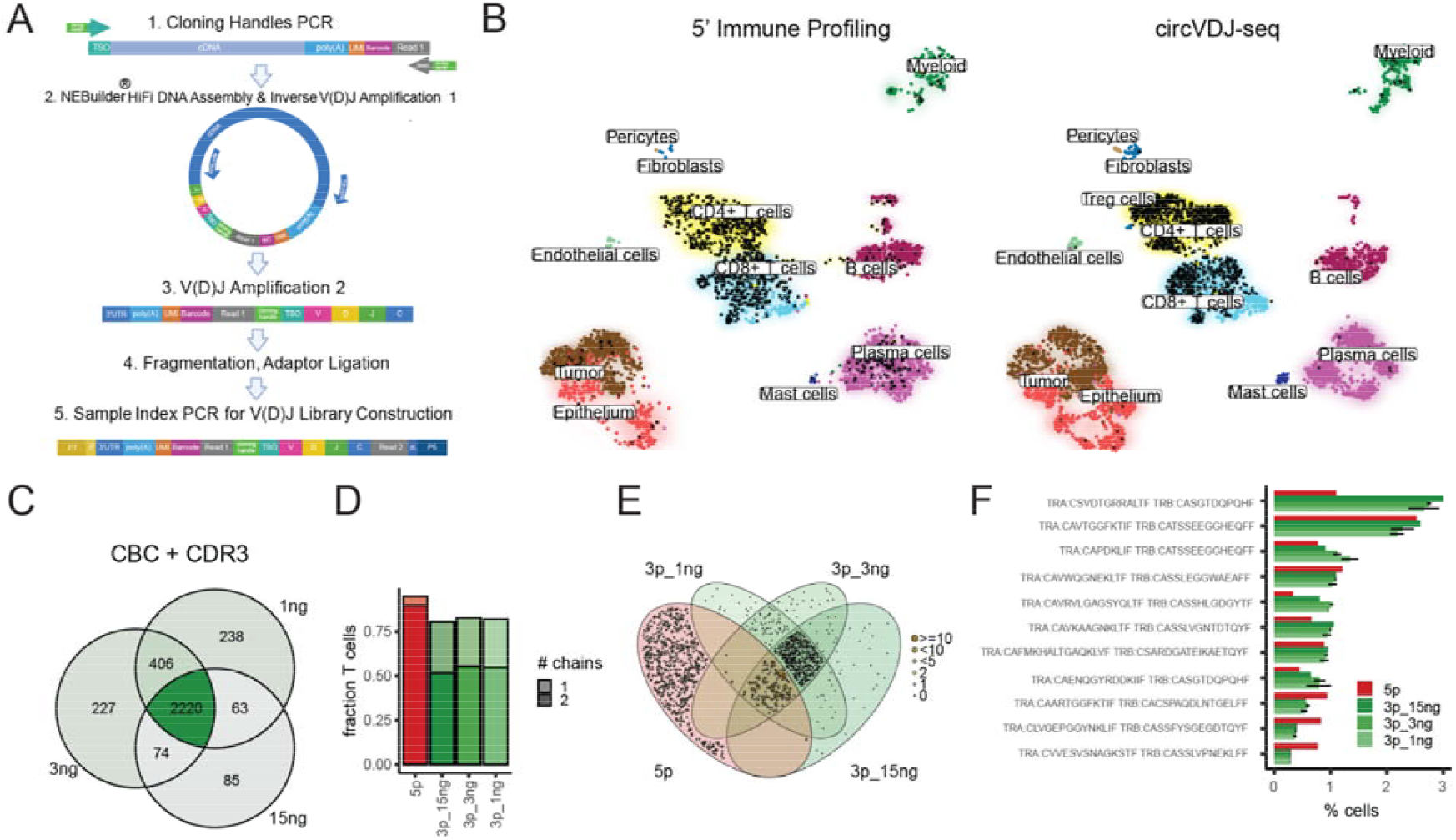
circVDJ-seq enables TCR profiling with 3’-directed single-cell RNA-seq. A) circVDJ-seq workflow: both ends of the cDNA library are appended by short homologous sequences to enable circularization via Gibson assembly and reposition 3’ cell barcodes next to the 5’end of cDNA molecules. This is followed by TCR-specific cDNA amplification, fragmentation, and short-read library amplification. B) UMAP plot showing gene expression data from CRC samples with identified TCR clones highlighted as black dots. C) overlap of identified combinations of CDR3 sequences and cell barcodes in circVDJ-seq replicates derived from the same cDNA library at different input concentrations. D) Fraction of T cells containing one or two TCR chains in the 5’IPv2 assay vs. circVDJ-seq at different input concentrations. E) Venn diagram shows overlap between the clonotypes recovered by the different libraries. Sizes of depicted clones are based on the frequencies observed with 5’IPv2. F) TCR clonotype frequencies. Error bars from SEM across two technical circVDJ-seq replicates.

To validate the approach, we made use of archived cDNA from a microsatellite-unstable colorectal cancer sample that we previously processed in parallel with the 3’GEX v3.1 and 5’IPv2 assays to investigate signaling pathways in oncogenic cell trajectories (**Fig. 1B**) [14,20]. Increasing amounts of 3’GEX cDNA were used as input for circVDJ-seq library preparation, resulting in comparable performance of the assay. Since the circVDJ-seq read structure mirrors the outputs obtained from the 5’IPv2/3 assay, we were able to readily process circVDJ-seq data with the VDJ Cell Ranger pipeline after replacing the cell barcode (CBC) whitelist with barcodes from the 3’GEX v3.1 assay. The resulting output data were further processed with Dandelion for improved VDJ contig annotation [25]. While circVDJ-seq data can be processed with other tools, combining Cell Ranger and Dandelion resulted in better reproducibility of technical replicates than processing the data with Cell Ranger alone, MiXCR [23], or TRUST4 [24], after pooling replicates and downsampling to the same number of input reads (**Additional file 2: Fig. S1C, D**). On average, 85% of circVDJ-seq sequence reads contained a valid CBC, and 65% of reads mapped to a VDJ gene (**Additional file 1: Table S3**). The detected CBCs, CDR3 sequences, and CBC+CDR3 combinations strongly overlapped between circVDJ-seq replicates, independently of the amount of input cDNA (**Fig. 1C**). On average, 68% and 90% of T cells contained a TCRα or TCRβ contig, yielding 59% of T cells with paired clonotype information, with highly similar sensitivity for different cDNA inputs (**Fig. 1D**), and a similar relationship between sequencing depth and clonotype recovery as observed for the 5’IPv2 assay (**Additional file 2: Fig. S1E**). Clonotypes determined by circVDJ-seq replicates were also readily recovered in relevant T cell clusters in 3’ gene expression data [17] (**Fig. 1B**).

Consistent with earlier studies of T cell receptor repertoires, we observed that the same TCRβ chain can be paired with different TCRα chains in the same individual [4,28], and that two TCRα chains can be co-expressed in the same cell [29,30]. Since these instances were relatively limited in our data set (**Additional file 2: Fig. S1F**), we decided to impute missing TCRα or TCRβ chains from the most frequent pairing to enable a better comparison of clonotype frequencies between replicates and assays. Individual clonotype frequencies appeared highly consistent across circVDJ-seq replicates and were largely independent of cDNA input amounts (**Fig. 1E, F**), suggesting that circVDJ-seq can be retrospectively applied even in scenarios where the available amount of archived cDNA is limited. While 5’IPv2 is expected to detect a larger overall number of TCR molecules due to the more direct assay design (**Fig. 1D, Additional file 1: Table S3**), the same amplified clonotypes were recovered by 3’ circVDJ-seq in a distinct set of cells from the same tumor, further supporting the accuracy of the approach (**Fig.1 E, F**).

### circVDJ-seq reliably detects TCR clonotypes from single cells and nuclei in multi-omics assays

Linking TCR sequence and gene expression states with information about accessible chromatin regions in the same cells could provide crucial insights into the epigenetic regulation of T cell development and function. We therefore wanted to test if circVDJ-seq can recover TCR clonotypes from the ATAC+RNA Multiome assay (MO), which uses isolated nuclei as input material. For accurate benchmarking, we processed PBMCs from the same healthy donor with the 5’IPv2, 3’GEX v3.1 and MO assays in parallel. As before, 3’GEX v3.1 and MO circVDJ-seq sequence data were processed with Cell Ranger updated for the respective CBC whitelist, followed by Dandelion [25]. Mapped reads contained approximately 94% and 85% valid CBCs, approximately 89% and 74% could be mapped to a VDJ gene, (**Additional file 1: Table S3**), and the resulting CBC+CDR3 combinations showed very high overlap in triplicate circVDJ-seq libraries (**Additional file 2: Fig. S2A**). Upon direct comparison, 5’IPv2 and circVDJ-seq from 3’GEX v3.1 efficiently recovered T cell clonotype information, with circVDJ-seq nearly matching overall clonotype recovery of the 5’IPv2 kit (87% vs 97%) (**Fig. 2A**). At the same time, the number of T cells with assigned clonotypes was reduced to 36% in the MO assay, where we also obtained fewer cells with paired clonotype information due to a lower recovery of TCRα contigs, most likely caused by reduced transcript abundance in single nuclei compared to whole cells (**Fig. 2A, Additional file 2: Fig. S2B**). Interestingly, while TCR gene expression UMI counts are also lower in 5’IPv2 compared to 3’GEX v3.1 data, the more direct capture of VDJ sequences at the 5’-end of transcripts still ensures high VDJ sensitivity of the 5’ assay (**Additional file 2: Fig. S2B**).

**Fig. 2.**
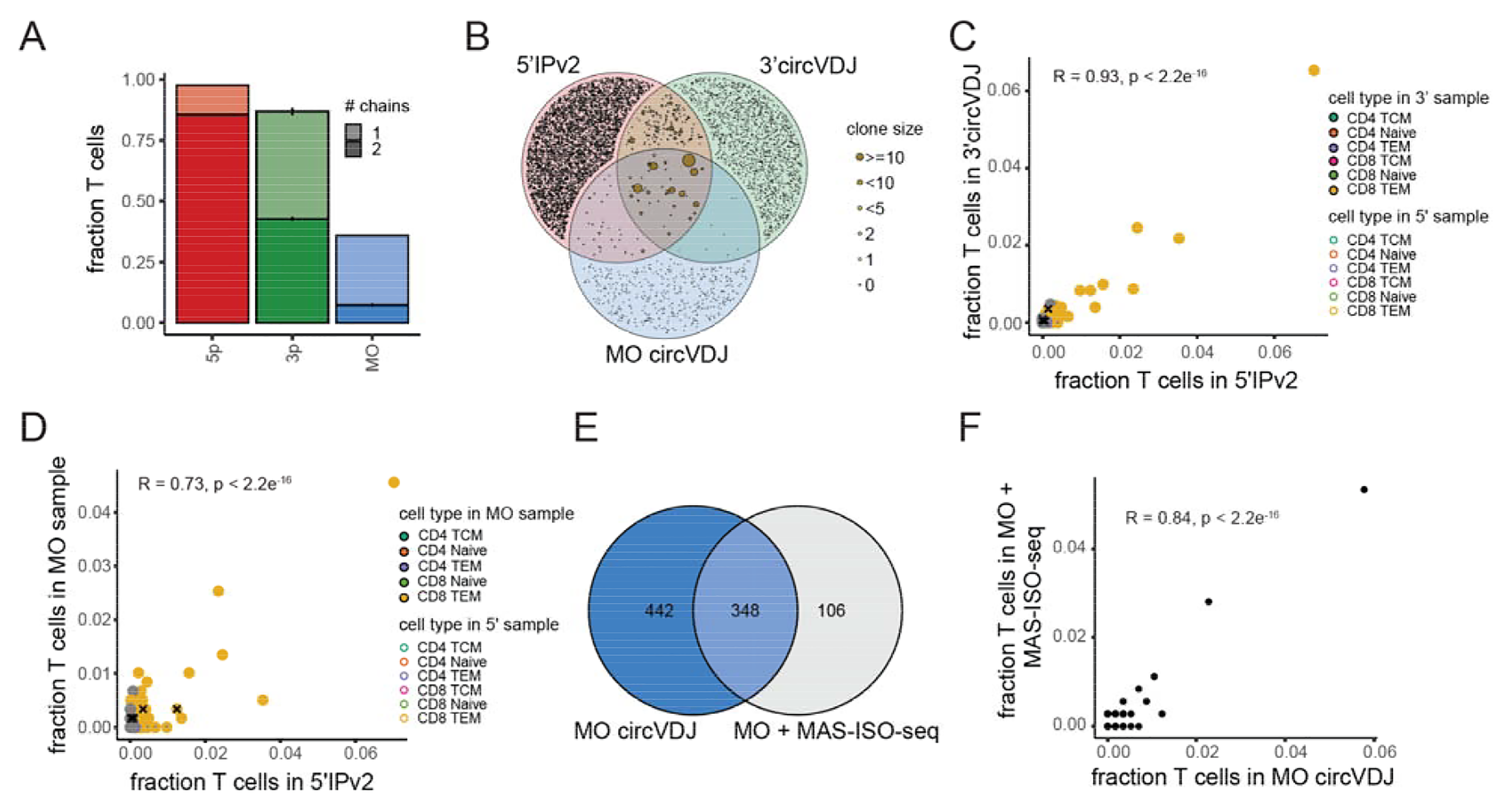
Robust TCR clonotype identification across single-cell multi-omics workflows. A) Fraction of T cells with assigned clonotypes in 5’IPv2, 3’ circVDJ-seq and MO circVDJ-seq. Error bars (for 3’ and MO assays) from SEM across 3 technical replicates. B) Venn diagram shows overlap between the clonotypes recovered by all three methods. Sizes of overlapping clones are based on the frequencies observed with 5’IPv2. C) Scatter plot shows the frequencies of individual T cell clones detected with 3’ circVDJ-seq or 5’IPv2 in PBMCs from the same donor. Fill and border color indicate the cell types assigned to each clone by the respective matched gene expression library, and crosses mark clones that are assigned to differing cell types in both assays. D) Scatter plot shows clonotype frequencies detected with MO circVDJ-seq and 5’IPv2 in PBMCs from the same donor as in C). E) Number of T cell clones detected by MO circVDJ-seq and MAS-ISO long-read sequencing of the same MO cDNA. F) Scatter plot shows T cell clonotype frequencies detected by MO circVDJ-seq and MAS-ISO-seq of the same MO cDNA.

We again imputed missing TCRα or TCRβ chains from the most frequent pairing to facilitate the comparison of clonotype frequencies between assays (**Additional file 2: Fig. S2C**). Although distinct populations of cells from the same donor were processed in each experiment, 4% of all clonotypes identified by 5’IPv2 were also found by circVDJ-seq from 3’-directed scRNA-seq after merging technical replicates, and 1.5% were found by circVDJ-seq from the MO assay (**Fig. 2B**). However, these recurring clonotypes represented 34% and 26% of cells, as 19 and 15 of the top 20 most abundant clones detected in the 5’IPv2 workflow were also recovered by circVDJ-seq from 3’GEX or MO cDNA, respectively. Consequently, the relative abundance of individual clones was highly correlated between circVDJ and 5’IPv2 (*R* = 0.93, *P* < 2.2e^-16^ and *R* = 0.78, *P* < 2.2e^-16^, respectively; **Fig. 2C, D; Additional file 2: Fig. S2D**), and the associated gene expression data reproducibly identified expanded clones as CD8 effector memory T cells. Interestingly, clonotype detection by circVDJ-seq showed similar efficiency in two unrelated PBMC samples that underwent a mild paraformaldehyde (PFA) fixation prior to MO processing as used in DOGMA-seq [31], highlighting the potential for parallel high-throughput TCR profiling alongside four separate genomic modalities at the single-cell level (**Additional file 2: Fig. S2E**).

Slight variations in individual clonotype abundances observed between MO circVDJ-seq and 5’IPv2 measurements from the same donor could be caused to varying degrees by fluctuations during multiple sub-sampling of cells, differences in the source material (whole cells versus nuclei), or by inaccuracy of the circVDJ-seq workflow itself. To exclude the latter possibility, we next validated the results from MO circVDJ-seq using an alternative approach. To this end, we directly subjected MO cDNA to long-read sequencing using MAS-ISO-seq without prior enrichment or amplification of VDJ sequences [32]. The overall number of detected clonotypes was reduced in MAS-ISO-seq compared to circVDJ-seq, which was expected since circVDJ-seq is a targeted approach and PacBio sequencing has limited sequencing depth (**Fig. 2E**). Importantly, however, clonotype abundances were highly correlated (*R* = 0.84, *P* < 2.2e^-16^) between both methods (**Fig. 2F, Additional file 2: Fig. S2D, F)**. Together, these data indicate that circVDJ-seq can faithfully recover TCR sequence information from 3’-barcoded cDNA libraries generated from whole cells, or from workflows that use single nuclei as starting material, such as the MO assay.

### Spatial circVDJ-seq detects T cell clonal expansion in autopsy-derived tissue

We next explored the utility of circVDJ-seq for spatial T cell clonotype mapping in autopsy-derived tissue samples. We recently identified a *CCL21* to *CCR7* signaling axis linked to the accumulation of exhausted T cells in ectopic lymphoid structures during prolonged lung immunopathology in COVID-19 by applying Visium gene expression profiling to lung and lymph node samples from deceased patients [13]. We generated triplicate circVDJ-seq libraries from Visium cDNA of a chronic COVID-19 postmortem lung sample from the same cohort that showed prominent immune cell infiltration and fibrotic areas with exacerbated collagen expression. We processed circVDJ-seq sequence reads with Cell Ranger VDJ after exchanging the CBC whitelist with Visium spot barcodes. Interestingly, although overall mapping statistics appeared similar to data from single-cells or nuclei (**Additional file 1: Table S3**), Cell Ranger VDJ did not report any spots with a successfully reconstructed TCR chain. This was potentially caused by the lower number of spot barcodes compared to single-cell barcodes, which might interfere with effective UMI thresholding. However, further processing of the raw Cell Ranger output with Dandelion [25] efficiently reconstructed TCRα and TCRβ chains together with their spatial addresses **(Fig. 3A, B)**.

**Fig. 3.**
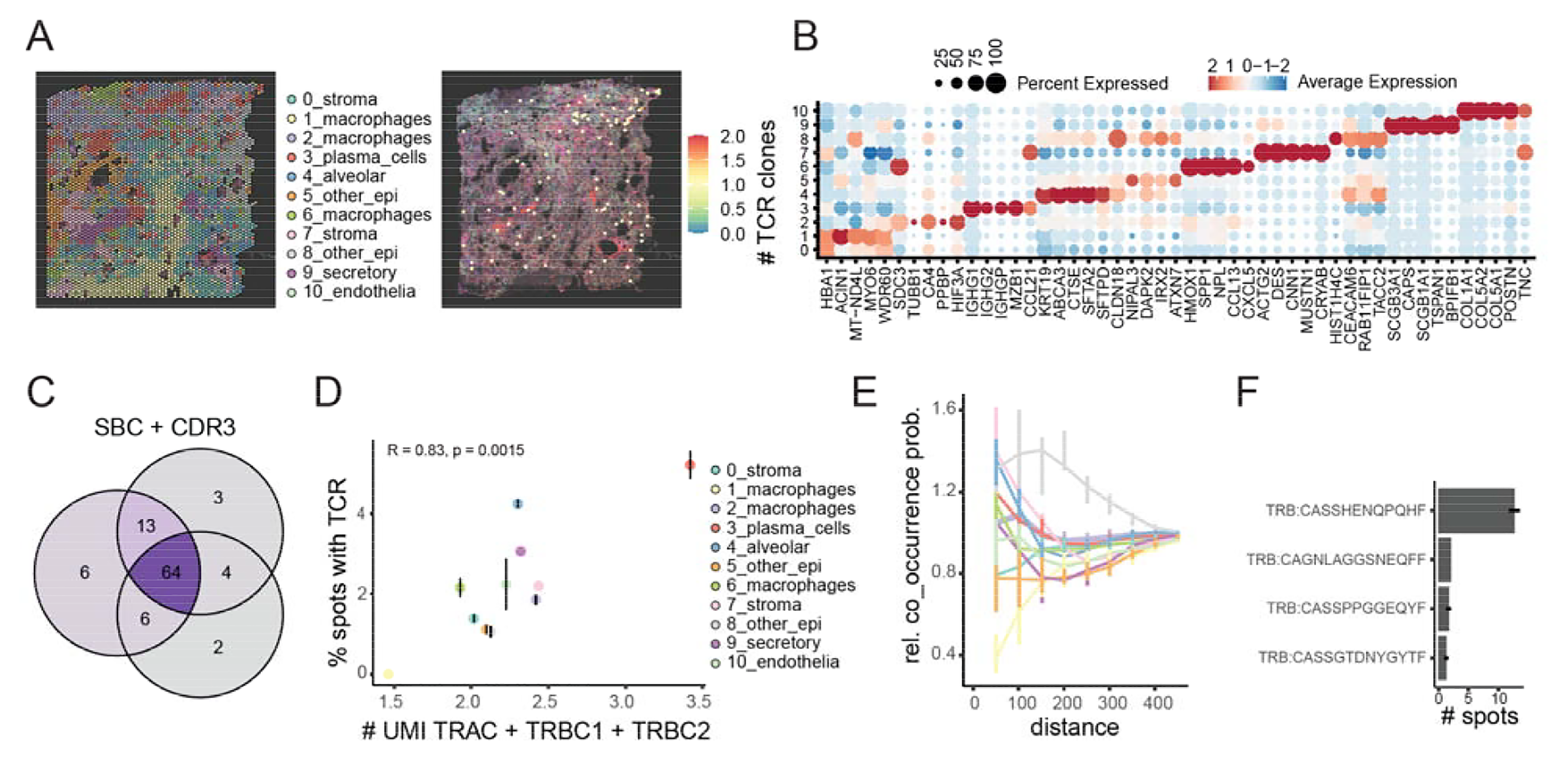
Spatial circVDJ-seq reveals clonal T cell expansion in autopsy-derived COVID-19 lung tissue. A) Gene expression clusters and TCR clones identified by Visium and circVDJ-seq in chronic COVID-19 lung (chronic case 5). B) Marker genes for clusters shown in A. C) Venn diagram shows overlap between the SBC and CDR3 combinations detected in technical circVDJ-seq replicates. D) Correlation between Visium TCR gene expression UMI counts and the fraction of spots with circVDJ-seq TCR clones found in the same clusters. Error bars indicate SEM across technical triplicates. E) Relative co-occurrence probabilities of spots with TCR clonotypes with spots from other clusters within a certain distance. Error bars indicate std. deviation across random subsamples of index spots. F) Reproducible detection of identified TCR clonotypes, error bars correspond to three circVDJ-seq replicates. Error bars indicate SEM across technical triplicates.

The identified T cell clonotypes were highly concordant between technical replicates (**Fig. 3C**), and the proportion of spots with detected TCR clones per cluster correlated with TCR UMI counts in the corresponding Visium GEX data (**Fig. 3D**). This analysis revealed the highest T cell abundance in plasma cell and alveolar spots, and in close proximity to stroma cluster 7 and cluster 8 spots (**Fig. 3D, E**), as calculated by relative co-occurrence probability between spots with TCR and spots of other clusters within a certain radius [26]. circVDJ-seq further revealed substantial expansion of a single clonotype with TCRβ CDR3 amino acid sequence CASSHENQPQHF (**Fig. 3F**). We previously observed T cell activation markers in COVID-19 lung-draining lymph nodes (dLNs) with no signs of active viral infection, suggesting that T cell activation persists in dLNs after the infection has been resolved [13]. We therefore expanded our analysis to archived Visium cDNA from an acute, chronic, and prolonged COVID-19, and a non-COVID-19 pneumonia dLN from the same cohort [13], to determine the respective extent of clonal T cell expansion (**Fig. 4A, B**). To further benchmark sensitivity and accuracy of circVDJ-seq, we additionally profiled the acute and chronic COVID-19 as well as the non-COVID-19 pneumonia LN with the recently introduced Spatial VDJ method [8]. Spatial VDJ is based on pre-enrichment of TCR sequences via a panel of capture oligos, which is followed by long-read sequencing (LR Spatial VDJ), or by multiplexed PCR against all V genes and short-read sequencing (SR Spatial VDJ) [8]. We chose LR spatial VDJ for comparison with circVDJ-seq, since the authors reported higher sensitivity towards TCR sequences compared to SR Spatial VDJ [8]. The detected CDR3 to spot-barcode combinations and clonotype frequencies appeared remarkably concordant between both methods (**Fig. 4C, Additional file 2: Fig. S3**).

**Fig. 4.**
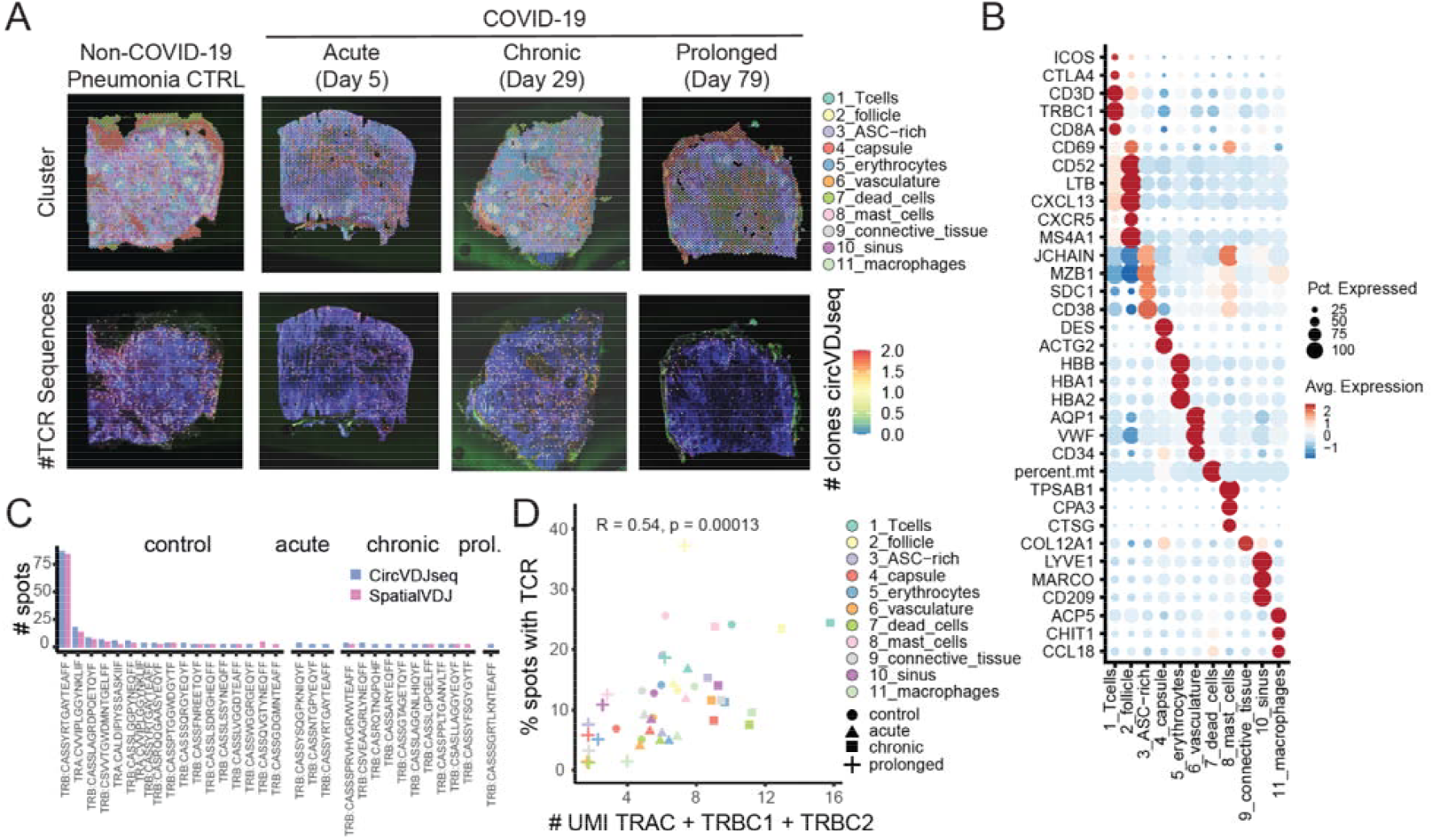
Spatial circVDJ-seq shows distinct clonal T cell expansion in lung-draining lymph nodes. A) Visium gene expression clustering and identified TCR clones in autopsy-derived lymph nodes from non-COVID-19 pneumonia, and acute, chronic, and prolonged COVID-19. B) Marker genes for gene expression clusters shown in A. C) TCR clonotype frequencies for clones identified in >2 spots for circVDJ-seq and LR Spatial VDJ. D) Correlation between TCR gene expression UMI counts and percentage of spots with identified TCR clones for individual gene expression clusters.

At the same time, circVDJ-seq consistently recovered more TRA and TRB chains compared to LR spatial VDJ, indicating at least matching sensitivity compared to the more resource-intensive assay (**Additional file 2: Fig. S4**). Further analysis of circVDJ-seq data showed that the frequency of TCR detection again strongly correlated with the number of TCR gene expression UMIs in the same gene expression clusters (**Fig. 4D**), and identified TCR clones were most abundant in the T cell zone surrounding B cell follicles (**Fig. 4B, D**). At the same time, we observed very low levels of clonal expansion in COVID-19 LNs (**Fig. 4C**). In contrast, the non-COVID-19 control sample showed very pronounced clonal T cell expansion (**Fig. 4C**), potentially caused by non-COVID-19 pneumonia and a possible squamous cell carcinoma lymph node metastasis of this donor, as suggested by concomitant keratin expression. Together, these data indicated that circVDJ-seq can efficiently retrieve contrasting clonotype dynamics even from challenging samples such as autopsy-derived tissue.

### Spatial circVDJ-seq reveals differential T cell infiltration in high- and low-risk neuroblastoma

To further explore its clinical utility, we finally applied circVDJ-seq to freshly resected neuroblastoma samples. Neuroblastomas are neural crest cell-derived pediatric tumors and represent the third most common type of cancer in children. NB cases can be categorized into high-, intermediate-, and low-risk disease based on patient age, histology, and genetic alterations like DNA ploidy and *MYCN* amplification [33]. Stage 4s neuroblastomas constitute a special group of low-risk tumors that are disseminated but often spontaneously regress. While patients with low-risk disease have a very good prognosis, the 5-year survival rate of high-risk patients remains at ∼50%. To test if circVDJ-seq can detect differences in the immune tumor microenvironment between distinct NB risk groups, we first generated Visium libraries from a stage IV high-risk (HR) and a stage 4s low-risk (LR) neuroblastoma, both without *MYCN* amplification.

Joint analysis of spatial gene expression revealed four tumor cell clusters and five stroma cell clusters with differential enrichment in the LR- and HR-NB sample (**Fig. 5A-C**). LR-NB tumor clusters 0, 1 and 3 showed high expression of the known low-risk markers *PRPH* and *NRCAM* [34,35] and several genes associated with favorable prognosis were additionally highly expressed in LR-NB cluster 3, including *CCNL2, WSB1, PCBP4*, and *HAND2-AS1* [36–41]. In contrast, the predominant HR-NB tumor cluster 2 showed downregulation of neuronal differentiation markers, low levels of *PLXNA4* and high expression of sympathetic *NPY* and *NEAT1*, which have all been linked to poor prognosis [42–45].

**Fig. 5.**
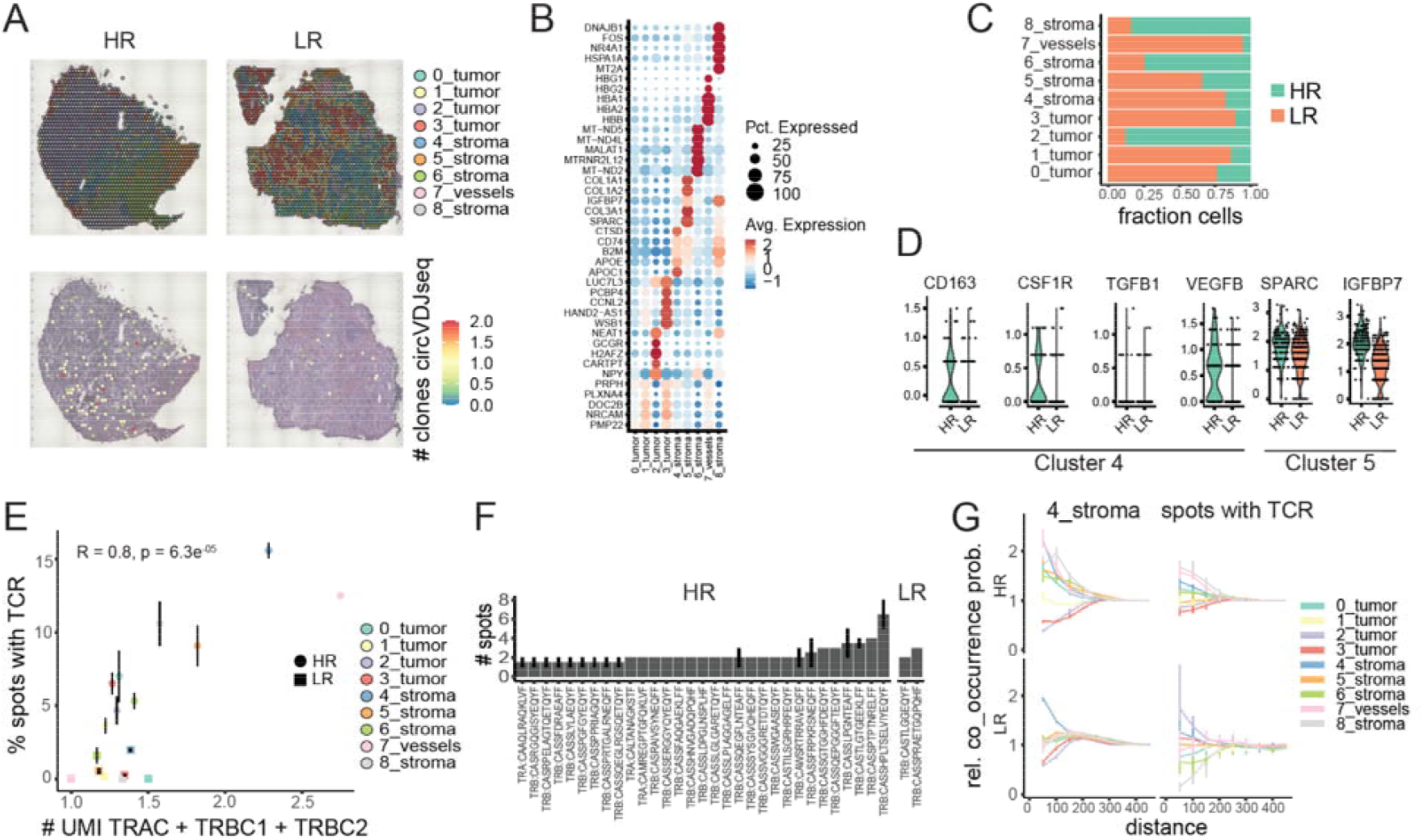
Spatial circVDJ-seq reveals distinct T cell dynamics in HR and LR neuroblastoma. A) Gene expression clusters (top) and TCR clones (bottom) identified by Visium and circVDJ-seq of high-risk (HR) and low-risk (LR) neuroblastoma. B) Marker genes for clusters shown in A. C) Cluster composition of HR-NB and LR-NB sample for the clusters in A. D) Expression of *CD163, CSF1R, TGFB1* and *VEGFB* in cluster 4, and *SPARC* and *IGFBP7* in cluster 5 in HR and LR neuroblastoma. E) Correlation between Visium TCR gene expression UMI counts and the fraction of spots with a circVDJ-seq TCR clone in individual gene expression clusters. Error bars indicate SEM across technical duplicates. F) Frequencies of spatially expanded clonotypes in the HR and LR neuroblastoma sample. Error bars indicate SEM across technical duplicates. G) Relative co-occurrence probabilities of spots from cluster 4 (left panel) and of spots with TCR clonotypes (right panel) with spots from other clusters within a certain distance in HR and LR neuroblastoma. Error bars indicate std. deviation across random subsamples of index spots.

Besides differences in tumor cell states, both samples also displayed substantial differences in tumor microenvironment (TME) composition. LR-NB showed higher abundance of cluster 4 with high expression of *CD74, APOE* and *APOC1*, reflecting immune cell infiltration [46,47]. At the same time, immune cluster 4 spots in HR-NB showed higher expression of *CD163, TGFB1, CSF1R* and *VEGFB*, indicating the presence of immunosuppressive tumor-associated M2 macrophages (**Fig. 5D**). Stromal cluster 5 was found in both LR- and HR-NB and contained markers of cancer-associated fibroblasts (CAFs) *SPARC* and *IGFBP7* [48–51], but with higher expression of the CAF-secreted extracellular matrix (ECM) glycoprotein *IGFBP7* in HR-NB (**Fig. 5D**). Stromal clusters 6 and 8 were strongly enriched in the HR-NB sample. Myeloid markers and higher levels of mitochondrial genes in cluster 6 might point towards metabolic stress of intratumoral myeloid cells [52], while cluster 8 showed signs of stress response and inflammation in a reactive stroma and high levels of *MT2A* indicative of CAFs linked to disease progression and poor prognosis [53,54]. Taken together, this analysis revealed substantial differences in tumor cell state and microenvironment, with a less aggressive cancer state and enhanced immune infiltration in LR-NB, and signs of immunosuppression, inflammatory stress and CAFs in the HR-NB patient sample.

We next generated duplicate circVDJ-seq libraries from each sample and processed the sequenced reads with Cell Ranger VDJ and Dandelion. This analysis revealed vastly different levels of T cell clonotypes in the HR- and LR-NB samples with 116 and 16 identified T cell clones, respectively (**Fig. 5A; F**). The number of spots that contain T cell clones in HR-NB strongly correlated with TCRαβ UMI counts in individual gene expression clusters (**Fig. 5E**), showing the highest enrichment in blood vessels and immune cluster 4. circVDJ-seq revealed substantial differences in cell type composition of immune cluster 4 between the LR-NB and HR-NB patient sample, with strongly elevated T cell counts and clonal amplification in HR-NB compared to low T cell counts in LR-NB (**Fig. 5E, F**). In the HR-NB sample, cluster 4 spots mostly co-occurred with blood vessels, other immune cluster 4 spots, and stromal spots, but were mostly excluded from the vicinity of the predominant HR-NB tumor cluster 2 (**Fig. 5G, top left panel**). The same exclusion from the HR-NB tumor area was also observed when the analysis was centered on all spots that include a T cell clone (**Fig. 5G, top right panel**). Conversely, in the LR-NB sample immune cluster 4 spots were not excluded from the LR-enriched tumor clusters 0 and 1, but appeared more distant from LR-NB tumor cluster 3. However, centering the analysis on the few spots that contain a T cell clone showed no spatial exclusion from the tumor area in general. Taken together, these analyses indicated substantial differences in the immune microenvironments of the HR- and LR-NB sample, with a more pronounced myeloid immune infiltration in LR-NB, and a higher abundance of clonally expanded cells in HR-NB, which remained largely excluded from the tumor area and were accompanied by immunosuppressive M2 macrophages and signs of a stressed reactive stroma.

## Discussion

TCR profiling provides essential insights into adaptive immunity, informing diagnosis, prognosis, and therapeutic approaches in cancer, infection, and autoimmune diseases. Here, we introduced circVDJ-seq, a streamlined and cost-efficient method for comprehensive TCR clonotype analysis from widely used 3’-barcoded transcriptomics assays. Unlike existing methods requiring complex primer sets or costly sequencing technologies, circVDJ-seq simplifies clonotype retrieval through cDNA circularization and amplification of conserved TCR constant regions. This reduces reagent and sequencing costs, but more importantly also eliminates inherent biases of the multiplexed PCR reaction that may lead to uneven amplification of different TCR clones [55]. In addition, circVDJ-seq can be applied to species with inaccurate or incomplete genome annotations of the TCR variable genes.

We show that circVDJ-seq accurately recovers TCR sequence information encoded in the cDNA libraries that serve as input material, evidenced by highly correlated results from technical circVDJ-seq replicates, parallel long-read sequencing of the original cDNA template, and 5’-directed TCR profiling of cells from the same donor. Although circVDJ-seq recovers clonotypes efficiently, sensitivity depends heavily on the quality of the initial cDNA libraries, emphasizing the importance of optimal sample preparation protocols. Besides timely acquisition and processing of biological samples, the underlying cDNA amplification method must amplify full-length cDNA molecules, for instance via a template switch mechanism, whereas workflows that use random hexamers for second-strand priming lead to shortened cDNA molecules that lack complete VDJ information.

While a 5’-directed strategy will provide the highest sensitivity for paired clonotype identification due to fewer library preparation steps, circVDJ-seq can efficiently reconstruct the clonotype composition of archived cDNA generated with 3’-directed scRNA-seq protocols, or from inherently 3’-directed assays such as the ATAC+RNA MO workflow. Our results from MO circVDJ-seq also show that TCR clonotype information can be retrieved from single nuclei. This enables the identification of the most abundant TCR clones, albeit with lower proportion of α chains and a reduced overall number of recovered clonotypes owing to the lower number of captured mRNA molecules in nuclei compared to whole cells. Nevertheless, single-nucleus RNA-seq has been the method of choice for a wide range of biological samples that are collected via snap freezing, e.g., during surgery, and circVDJ-seq enables the retrospective analysis of T cell clonality in such samples.

Similarly, circVDJ-seq can be combined with any spatial transcriptomics method that is based on poly(A) capture of mRNAs and template switching, such as Visium, Slide-seq [7,56] or Stereo-seq [57]. Direct comparison of circVDJ-seq with the recently introduced LR Spatial VDJ method revealed highly concordant clone frequencies between both methods, and at least equivalent sensitivity of circVDJ-seq, establishing circVDJ-seq as an easy to use and cost-efficient alternative to Spatial VDJ that does not require complex oligo panels or specialized sequencing equipment. Using the Visium workflow for initial cDNA library generation, we demonstrated that circVDJ-seq can provide important insights into T cell biology. Our spatial circVDJ-seq analysis identified distinct immune landscapes in neuroblastoma subtypes, with notable exclusion of T cells from tumor compartments in high-risk disease, consistent with the presence of immunosuppressive microenvironments previously documented in aggressive pediatric cancers. This observation is also consistent with recent spatial transcriptomics studies on high-risk neuroblastoma that have shown spatial compartmentalization with malignant tumor cells often spatially segregated from immune cells [52].

We furthermore observed enhanced clonal amplification in lung-draining lymph nodes from cancer-associated pneumonia compared to prolonged COVID-19, illustrating the utility of circVDJ-seq as a versatile tool for TCR profiling in various tissue contexts, including postmortem autopsy material, which was especially encouraging. Here, the integrity of the starting material is of particular importance, since degradation of mRNA molecules will render TCR information inaccessible. Timely tissue processing and the testing of RNA integrity are therefore advisable in this context. The examples shown here are based on standard Visium with 50 µm spot diameter, corresponding to multiple cells per spot. However, newer workflows such as STOMICS or Visium HD 3’ have reached true single-cell resolution combined with enhanced mRNA capture efficiency, and circVDJ-seq can be readily applied to those workflows with equivalent cDNA structures.

## Conclusions

circVDJ-seq is a simple and robust assay for TCR profiling from 3’-barcoded cDNA libraries. It is broadly applicable to single-cell multi-omics workflows as well as spatial transcriptomics and does not require large pools of PCR primers, capture oligos, or specialized long-read sequencing equipment, enabling easier access to single-cell and spatial TCR repertoire profiling. We envision that circVDJ-seq will facilitate widespread and cost-effective TCR profiling across diverse single-cell and spatial transcriptomics platforms and clinical contexts.

## Abbreviations

ATAC: assay for transposase-accessible chromatin
BSA: bovine serum albumin
CAF: cancer-associated fibroblast
CBC: cell barcode
cDNA: complementary DNA
CDR3: complementarity-determining region 3
circVDJ-seq: circularized VDJ sequencing
COVID-19: coronavirus disease 2019
CRC: colorectal cancer
DTT: dithiothreitol
dLN: draining lymph node
EB: elution buffer
ECM: extracellular matrix
GEX: gene expression
HD: high definition (in context of spatial transcriptomics)
HR-NB: high-risk neuroblastoma
IMGT: international immunogenetics information system
5’IPv2/3: immune profiling version 2/3
LR-NB: low-risk neuroblastoma
MAS-ISO-seq: multiplexed amplification of specific isoforms sequencing
MHC: major histocompatibility complex
MO: multiome (combined RNA and ATAC sequencing)
PBMC: peripheral blood mononuclear cell
PCA: principal component analysis
PCR: polymerase chain reaction
PFA: paraformaldehyde
qPCR: quantitative PCR
RNA: ribonucleic acid
RT-PCR: reverse-transcription PCR
SBC: spot barcode
scRNA-seq: single-cell RNA sequencing
SEM: standard error of mean
SPRI: solid-phase reversible immobilization
STOMICS: spatial transcriptomics omics
TCR: T cell receptor
TRAC: T cell receptor alpha constant region
TRBC: T cell receptor beta constant region
UMI: unique molecular identifier
UTR: untranslated region
VDJ: variable (V), diversity (D), joining (J)
UMAP: uniform manifold approximation and projection

## Declarations

### Ethics approval and consent to participate

This study was conducted in accordance with the Declaration of Helsinki and with the approval of the Ethics Committee of the Charité (EA2/066/20, EA1/317/20, EA2/144/15, EA1/005/21, and EA4/164/19), the Charité - BIH COVID-19 research board, and the Ethics Committee of the Medical Faculty of the University of Cologne (#9764 and #04-049). Autopsies were performed on the legal basis of §1 926 SRegG BE of the Autopsy Act of Berlin and §25(4) of the German Infection Protection Act. Written informed consent was obtained from the patients’ families prior to autopsy.

## Additional files

### Additional file 1

File format: Microsoft Word document (.docx)

Title of data: Supplementary Tables S1-S3

Description of data:

Table S1: Human samples used in this study

Table S2: Oligonucleotides used for circVDJ-seq

Table S3: Sequencing statistics

### Additional file 2

File format: PDF file (.pdf)

Title of data: Supplementary Figure S1, Supplementary Figure S2, Supplementary Figure S3,

Supplementary Figure S4,

Description of data:

Fig. S1: Amplified TCR cDNA library from a CRC sample visualized on TapeStation; TCR short-read sequencing library quality control; comparison of CBC+CDR3 combinations detected by different VDJ sequencing data-processing pipelines; relationship between sequencing depth and detected TCR clones; and frequency of TCRα or TCRβ chains detected in more than one pairing.

Fig. S2: Comparison of clonotype recovery and TCR detection across 5’ immune profiling, 3’ gene expression circVDJ-seq, multiome circVDJ-seq, and MAS-ISO-seq datasets from PBMCs, including replicate overlap, TCR UMI counts, clone sizes, and fraction of T cells with assigned clonotypes.

Fig. S3: Venn diagrams showing the overlap of detected CDR3 sequences, spatial barcodes, and CDR3-spatial barcode combinations between circVDJ-seq and long-read Spatial VDJ.

Fig. S4: Number of spatial barcodes with a detected TCR chain in circVDJ-seq and long-read Spatial VDJ.

### Consent for publication

Not applicable.

### Availability of data and materials

Processed data is available at https://zenodo.org/records/19336264 [58]. Analysis code and scripts to generate all figures are available at https://github.com/bihealth/circVDJseq_paper [59].

The raw sequencing data generated in this study from human tissue samples have been deposited in the German Human Genome-Phenome Archive (GHGA) under the accession code https://data.ghga.de/study/GHGAS15310500752602 [60]. These data are available under restricted access due to data privacy regulations and ethical requirements related to personal data from a vulnerable patient population. Access to the data is granted only to qualified researchers for non-commercial research use upon approval of a controlled access request. Requests for data access can be submitted via the GHGA Data Portal [https://data.ghga.de] or by contacting. Applications are reviewed by the Data Access Committee (DAC) at Charité - Universitätsmedizin Berlin in coordination with GHGA, and responses and data access are typically provided within 3–4 weeks. Approved applicants are required to sign a Data Use Agreement (DUA) governed by Charité - Universitätsmedizin Berlin. A copy of the DUA is available upon request from the corresponding author or from the DAC. Users of these data are requested to cite this study in any resulting publications. Data reuse is limited to non-commercial research purposes.

## Authors’ contributions

I.P. and T.C. conceived the circVDJseq workflow. I.P. performed circVDJseq experiments. B.O. developed circVDJseq analysis pipelines. Y.H., C.B., M.G., M.SZ., V.S. and C.D. generated single-cell RNAseq and Multiome data and provided cDNA for circVDJseq. I.T., T.M.P. and N.R. generated Neuroblastoma Visium data and provided cDNA for circVDJseq. A.PR. and A.E.H. generated lung and lymph node Visium data and provided cDNA for circVDJseq. C.Q., J.W. and T.B. performed long-and short-read sequencing. B.O., R.R. and C.F. performed data analysis. I.P., B.O., R.R., H.R., A.E.H, and T.C. interpreted the data. I.P., B.O., and R.R. prepared figure panels. T.C. wrote the manuscript. A.E.H., A.E., M.S., and H.R. contributed clinical specimens. L.S.L. and J.A. commented on the manuscript. All authors read and approved the final manuscript.

## Acknowledgements

We thank the patients who participated in this study. We thank Rogier Versteeg and Vedran Franke for critical reading of the manuscript.

