## Supplementary Figures for "circVDJ-seq for T cell clonotype detection in single-cell and spatial multi-omics"

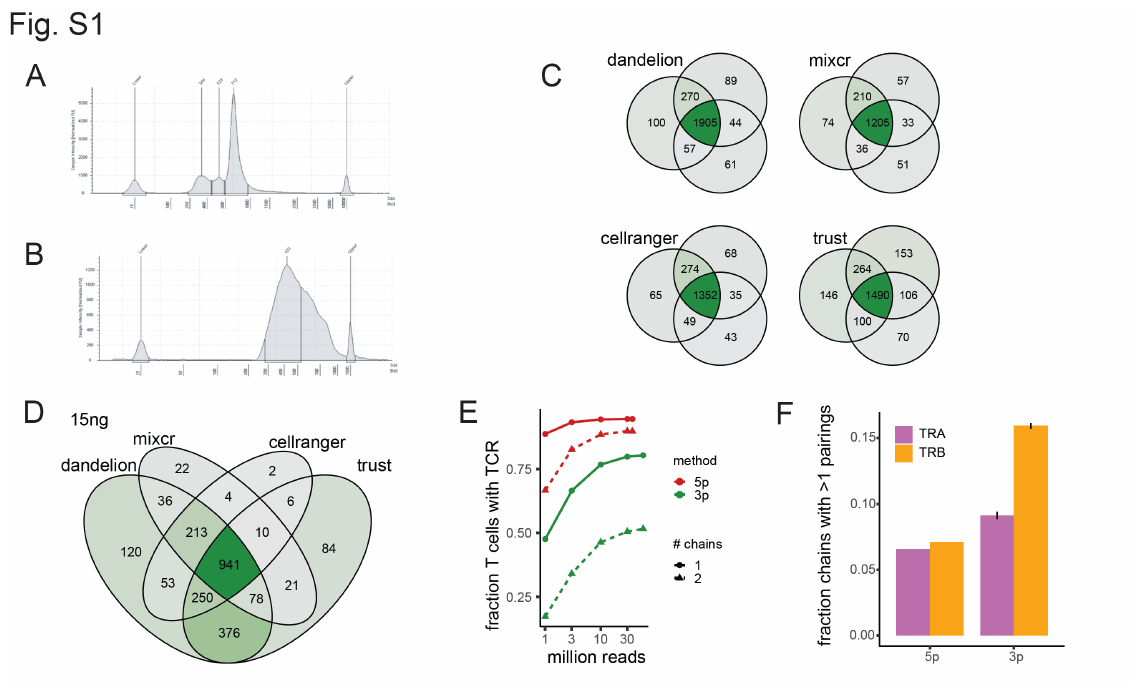

**Supplementary Figure 1**

Amplified TCR cDNA library from CRC sample visualized on Tapestation with HS5000 Screentape (Agilent). Initial cDNA was generated with the 3’GEX v3.1 assay (10X Genomics). B) TCR short-read sequencing library generated from (B) visualized on Tapestation with HS1000 Screentape (Agilent). C) Venn diagrams of CBC+CDR3 combinations detected with different VDJ sequencing data processing pipelines in circVDJ-seq samples from Fig. 1C, downsampled to identical numbers of input reads. D) Venn diagram of CBC+CDR3 combinations detected in the CRC 15ng circVDJ-seq sample with different processing methods. E) Relationship between sequencing depth and the number of detected TCR clones. F) Frequency of TCRα or TCRβ chains that are detected in more than one pairing in 5’IP v2 or 3’GEX v3.1 data. Error bars from s.e.m. across replicates.

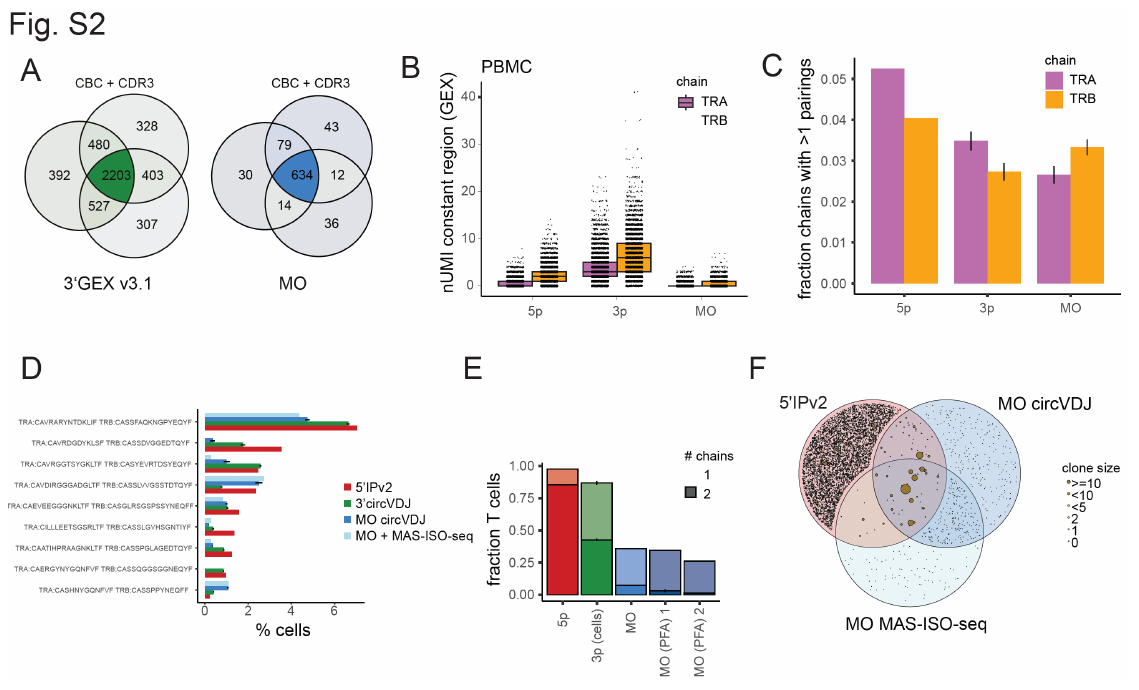

**Supplementary Figure 2**

A) Venn Diagram of the overlap of CBC+CDR3 combinations between circVDJ-seq replicates of 3’GEX v3.1 and ATAC+RNA MO data from PBMCs of the same donor. B) TCR UMI counts in 5’IPv2, 3’GEX v3.1 and MO GEX data from PBMCs of the same donor. C) Frequency of TCRα or TCRβ chains that are detected in more than one pairing in 5’IP v2 or circVDJ-seq data from 3’ or MO cDNA. Error bars from s.e.m. across technical replicates. D) Clone sizes for the top 10 TCRα+β clones (missing chains imputed) detected in 5’IP v2, circVDJ-seq from 3’ or MO cDNA, or MAS-ISO-seq data from MO cDNA, respectively. Error bars from s.e.m. across technical replicates. E) Fraction of T cells with assigned clonotypes in 5’IPv2, 3’v3.1 circVDJ-seq and MO circVDJ-seq as in Fig. 2A together with two additional MO datasets using mild PFA fixation. Error bars (for 3’ and MO assays) from s.e.m. across 3 technical replicates. F) Venn diagram as in Fig. 2B shows overlap between the clonotypes recovered by 5’ 5’IPv2 as well as circVDJ-seq and MAS-ISO-seq of the same MO cDNA. Sizes of overlapping clones are based on the frequencies observed with 5’IPv2.

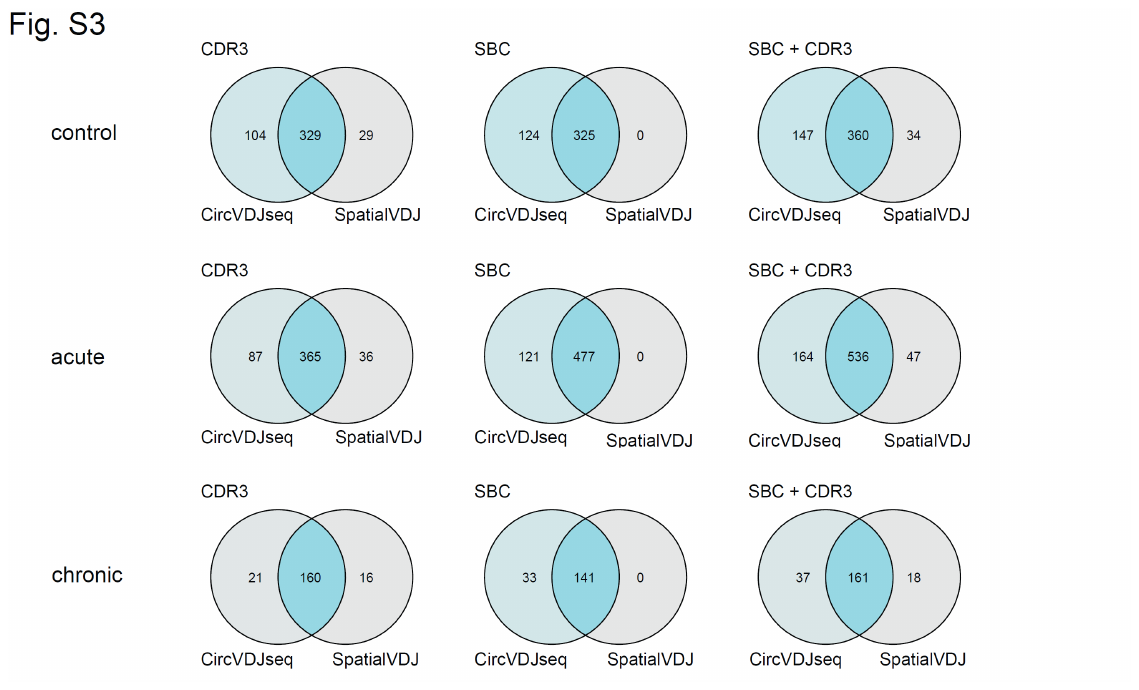

**Supplementary Figure 3**

Venn Diagrams showing the overlap of detected CDR3 sequences, spatial barcodes, and CDR3-SBC combinations between circVDJ-seq and LR Spatial VDJ.

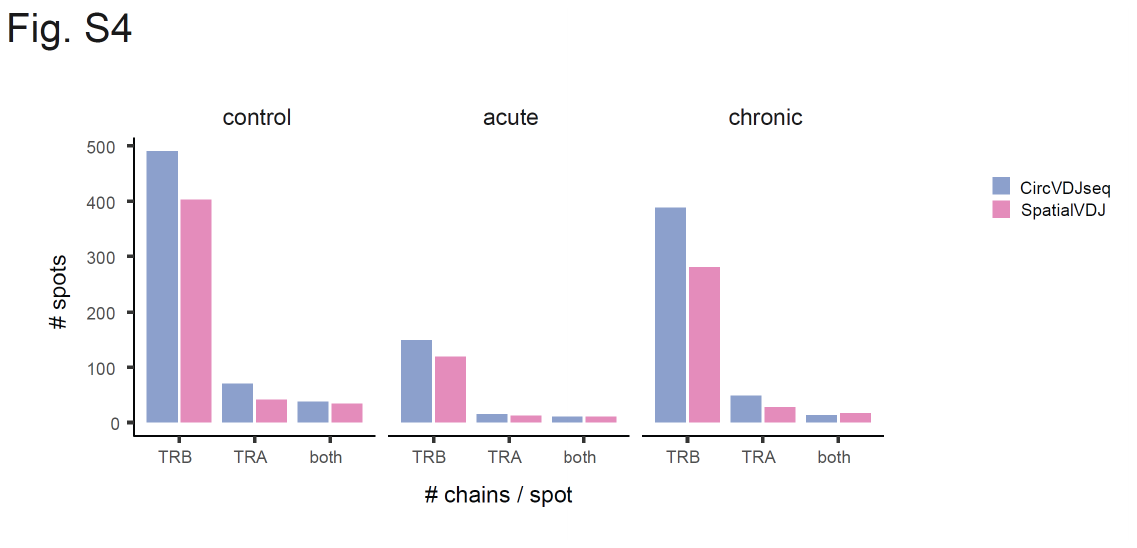

**Supplementary Figure 4**

Number of spatial barcodes with a detected TCR chain in circVDJ-seq and LR Spatial VDJ.
